# Rules for PP2A-controlled phosphosignalling and drug responses

**DOI:** 10.1101/271841

**Authors:** Otto Kauko, Susumu Y. Imanishi, Evgeny Kulesskiy, Teemu Daniel Laajala, Laxman Yetukuri, Anni Laine, Mikael Jumppanen, Pekka Haapaniemi, Luyao Ruan, Bhagwan Yadav, Veronika Suni, Taru Varila, Garry Corthals, Jüri Reimand, Krister Wennerberg, Tero Aittokallio, Jukka Westermarck

## Abstract

Systemic understanding of protein phosphatase 2A (PP2A)-regulated cellular processes is still at infancy. Here, we present mass-spectrometry analysis of phospho-targets (dephosphorylome) regulated by PP2A modulation. In addition to PP2A-regulated processes and targets, the data reveal important general concepts and rules related to PP2A-mediated phosphoregulation. These include the unidirectionality paradigm of regulation of phosphorylation, and differential spatial distribution of kinase-and phosphatase-dominated phosphotargets. Data also present first systemic analysis of targets of PP2A-modulating oncoproteins, CIP2A, PME-1, and SET; including targets via which PP2A may coordinately regulate activities of cancer drivers and tumor suppressors such as MYC or TP53. To validate functional utility of this dataset, PP2A dephosphorylome activity was correlated with cancer cell responses to over 300 drugs. Notably, we find that cancer therapy responses can be broadly classified based on PP2A dephosphorylome activity, both in quantitative and qualitative manner. In summary, our data characterize rules by which PP2A coordinate cancer cell phosphosignaling and drug responses. The results also may also direct the use of emerging pharmacological approaches for PP2A activity modulation in human diseases.

## Introduction

Serine/threonine phosphatase PP2A is essential for normal development^1^, and regulates large number of physiological and pathological processes ranging from immune responses^2^ to Alzheimer’s disease^3^. PP2A is also a critical human tumor suppressor, inhibition of which is a prerequisite for malignant transformation of many types of normal human cells ^4–7^. PP2A inhibition promotes *in vivo* tumorigenesis ^8–13^, and orally bioavailable, and non-toxic, PP2A reactivating small molecule compounds show robust antitumor effects ^14,15^. Although PP2A has been recognized as critical regulator of several signaling pathways, and RAS-driven oncogenic signaling ^6,9,16,17^, the depth by which PP2A controls cell signaling have not been systematically addressed as yet.

PP2A is a trimeric protein complex (Fig. 1A) in which a core dimer formed between the scaffolding A subunit (PPP2R1A, PPP2R1B) and the catalytic C subunit (PPP2CA, PPP2CB) is associated with one of the many B subunits that facilitate and direct the interaction of the trimer with substrate proteins ^16^. In addition to mutations found in cancer, PP2A activity is regulated by various non-genomic mechanisms^18^ including post-translational modifications of PP2A complex components, as well as by interaction with group of designated PP2A inhibitor proteins (PIPs hereafter). The best-characterized PIPs are CIP2A, PME-1, and SET (Fig. 1A)^4,19,20^. Despite their classification as PIPs, CIP2A, PME-1, and SET do not share structural features ^21–23^, and the mechanisms by which they inhibit PP2A activity toward their selected phosphoprotein targets are vastly different. SET binds and directly inhibits the catalytic center of PP2Ac ^24,25^, whereas PME-1 regulates PP2Ac activity via demethylation of the C-terminal leucine 309 and through the direct eviction of metal ions from the catalytic center ^20,21^. CIP2A binds the PP2A complex via a direct interaction with the B56 regulatory subunits ^23^. Importantly, despite their differential mode of PP2A regulation, the cancer-relevant phenotypes affected by CIP2A, PME-1 or SET modulation can be rescued by concomitant PP2A modulation ^10,26–28^, indicating that their function are truly dependent on PP2A. Tissue and cell type specific expression, and regulation of PIPs thus provides a sophisticated cellular mechanism to regulate the major Ser/Thr phosphatase PP2A. However, currently we lack any systematic understanding of processes regulated by, or functional redundancy of these PP2A modulating proteins. Also, the rules that determine functional consequences of PP2A modulation on cancer cellular signaling are very poorly understood.

**Fig 1.**
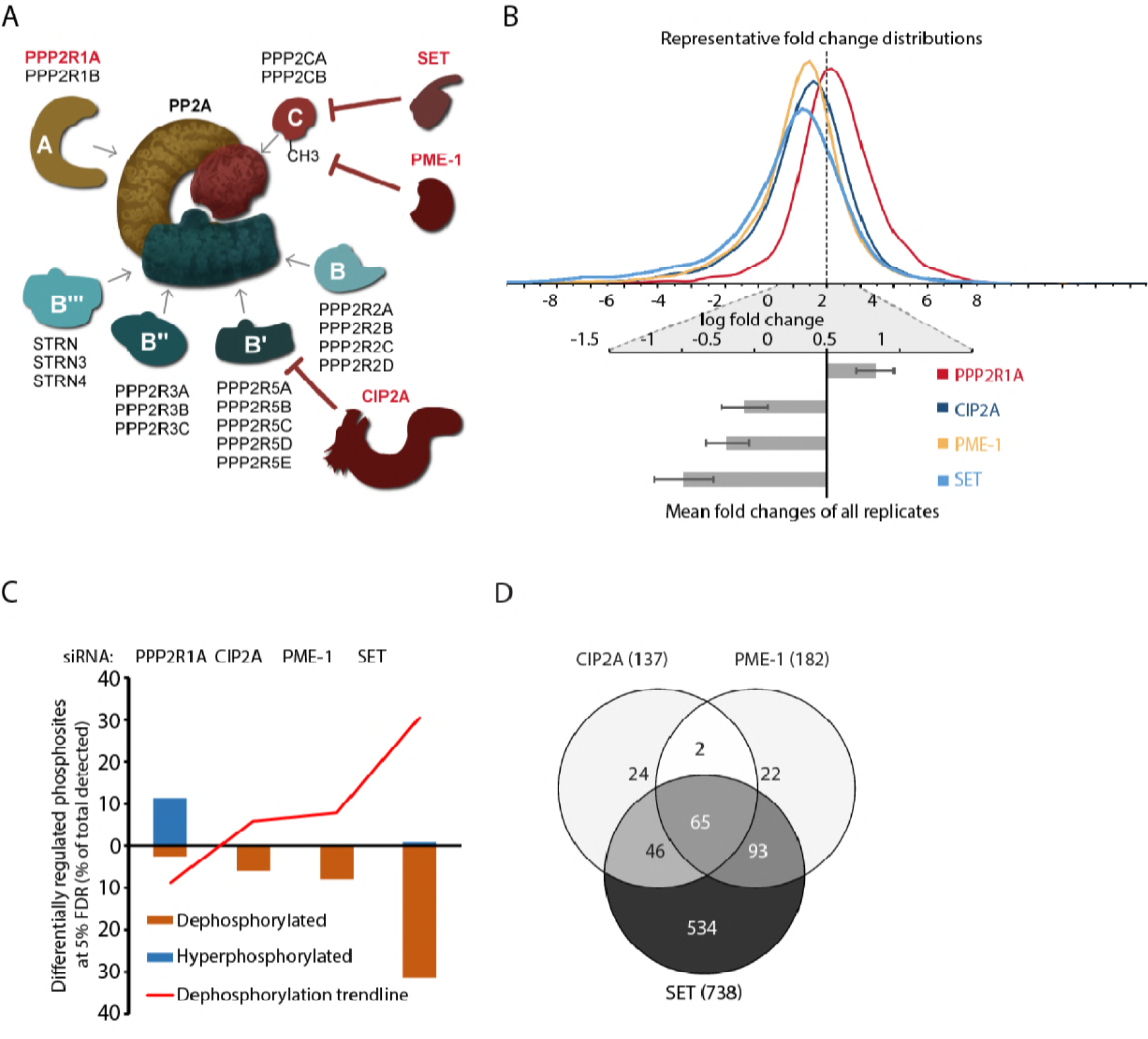
Identification and quantification of the PP2A dephosphorylome via LC-MS/MS. A) Schematic presentation of regulation of PP2A activity by subunit composition, endogenous inhibitor proteins (PIPs) and catalytic subunit methylation. B) Manipulation of PPPP2R1A or PIPs causes global shift in phosphorylation. Fold change distributions of representative replicates (top) and mean fold changes of all replicates (bottom). C) Fraction of differentially regulated phosphopeptides across all siRNAs for each gene (FDR <5%). PPP2R1A inhibition shows increased phosphorylation whereas inhibition of each CIP2A, PME-1 and SET caused almost exclusively protein dephosphorylation. D) Venn diagram showing the shared and unique differentially regulated phosphopeptides regulated by different PPI’s.

In concert with involvement of PP2A in tissue homeostasis, recent studies have revealed also specific roles for PIPs in a number of physiological and pathological processes. Physiologically CIP2A promotes spermatogenesis^29^, T-cell activation^30^ and intestinal regeneration in response to tissue damage^31^. PME-1 knock-out mice are perinatally lethal possibly due to problems with breathing and suckling behavior ^32^, and PME-1 overexpression has been linked to increased Tau phosphorylation and Alzheimer’s disease progression^33^. SET-mediated PP2A inhibition has instead been linked to natural killer cell activation ^34,35^. Notably, all these PIPs are commonly expressed at high levels in various cancer types, and inhibit PP2A phosphatase activity toward cancer-driving proteins. CIP2A in particular is a prevalently overexpressed oncoprotein across most human cancer types ^9,10,26,36^, and high CIP2A expression is a synergistic poor survival marker with RAS mutations and expression across TCGA pan-cancer data ^37^. Inhibition of these three PIPs also effectively suppresses malignant cell growth and tumorigenesis in *in vivo* models ^9,10,27,28^. Based on these characteristics, targeting of CIP2A, PME-1, and SET has recently emerged as an attractive novel therapeutic approach in cancers without PP2A mutations, particularly in combination with drugs that target phosphorylation-dependent signaling ^4,19,27^. However, whether the role for PIPs and PP2A in defining drug responses can be generalized beyond thus far tested kinase inhibitors is yet unclear.

Here we report phosphoproteome analysis of HeLa cells where we have targeted the PP2A scaffold protein PPP2R1A and CIP2A, PME-1, or SET. The study provides the largest currently available resource for understanding the processes targeted by PP2A activity. In addition to novel information on PP2A-regulated processes and targets, the data also reveal important general concepts related to phosphatase biology. These include the unidirectionality paradigm and differential spatial distribution of kinase-and phosphatase-dominated phosphotargets. An important implication of these findings is that deregulation of phosphatase activity alone can activate many established cancer relevant signalling pathways, particularly seen for MYC activity. Our data also demonstrate how PP2A dephosphorylome activity globally defines drug sensitivities. The insights provided by these results are important for holistic understanding of cell signaling but also may have translational relevance in light of emerging pharmacological approaches for PP2A activity modulation^14,15,38^ in human diseases ranging from Alzheimer’s disease to cancer.

## Results

### Identification and quantification of the PP2A dephosphorylome via LC-MS/MS

PPP2R1A is the predominant PP2A scaffold subunit, and essential for functional PP2A complex formation (Fig. 1A). Therefore, HeLa cells were treated with three independent PPP2R1A siRNAs to simulate PP2A inhibition. CIP2A, PME-1, and SET were targeted with independent siRNA sequences (3-4/gene) to characterize the phosphoproteomes that are regulated by three well characterized PIPs. HeLa cells were chosen as a model based on previous evidence of their robust but sub-lethal response to PP2A modulation ^26,37^.

Changes in peptide phosphorylation were analyzed 72 h after transfection via a previously described label-free phosphoproteomics method, combined with pairwise abundance normalization developed for accurate monitoring of global phosphorylation changes ^37^ (Fig. S1A). Importantly, the performance of the method was validated by high reproducibility between the phosphoproteomics data and phosphospecific antibody survey ^37^.

### Depletion of the PP2A scaffold PPP2R1A or PP2A inhibitor proteins results in global phosphorylation changes

In total, 7037 non-redundant phosphopeptides were quantified from the HeLa whole cell lysates. The reported PP2A dephosphorylome targets are limited to those that were quantified consistently across all replicate samples (Fig. S1D). Of the consistently quantified phosphopeptides, 43% were significantly regulated (5% FDR) by the selected manipulations, and 57% of the phosphoproteins had at least one differentially regulated phosphopeptide. The entire catalogue of differentially regulated phosphopeptides is included as Table S1 and will be available online at www.depod.org

Consistent with the role of PP2A as a potent serine/threonine phosphatase, depletion of the PP2A complex scaffold protein PPP2R1A resulted in a robust increase in the average phosphorylation level across the HeLa cell phosphoproteome (Fig. 1B, S1B-D). Notably, only a minor subset of the phosphopeptides significantly regulated by PPP2R1A exhibited decreased phosphorylation (Fig. 1C), which indicates that global PP2A inhibition cannot be acutely compensated for by the increased activities of other serine/threonine phosphatases.

In contrast, cells that were deficient in any of the PIPs (i.e., CIP2A, PME-1 or SET) (Fig. 1A) displayed global dephosphorylation (Fig. 1B, S1B-D). Depletion of either CIP2A or PME-1 caused similar overall increases in protein dephosphorylation, whereas SET clearly appeared to be the most robust inhibitor of protein dephosphorylation and significantly regulated 30.5% of the detected phosphopeptides (Fig. 1C). Importantly, because only a minor subset of the phosphopeptides that were significantly regulated by PIPs (0.5-2.7%) exhibited increased phosphorylation (Fig. 1C), these data also indicate that interference with serine/threonine dephosphorylation via SET, PME-1 and CIP2A inhibition cannot be acutely compensated for by increased kinase activities.

To address the outstanding question of the redundancy of CIP2A, PME-1 and SET in the regulation of global serine/threonine phosphorylation we assessed the degree of overlap between their dephosphorylome targets. In accord with a common regulatory mechanism of dephosphorylation inhibition (i.e., PP2A inhibition), 49% of the CIP2A targets were regulated also by PME-1, whereas 37% of PME-1-regulated targets were also CIP2A targets (Fig. 1D). The non-redundant targets could be due to the differential selectivity of CIP2A and PME-1 for different PP2A trimer complexes ^9,23,39^. On the other hand, SET clearly possessed the largest number of dephosphorylome targets; 534 targets were uniquely regulated by SET (Fig. 1D). The larger number of SET targets compared to CIP2A and PME-1 may be at least partly due to the direct inhibition of the catalytic activity of PP2Ac by SET ^24,25^ and the lack of selectivity towards specific B-subunit containing PP2A trimers, as is the case with PME-1 or CIP2A ^9,23,39^. Moreover, it should be noted that many SET targets were indeed regulated to certain extent also by CIP2A and PME-1 depletion (Table S1), although these effects did not meet the selected statistical filtering criteria. This overlap of targets regulated by PIPs, together with the tendency toward the enrichment of the recently identified PP2A B56 subunit binding motif ^40^ among the most significantly regulated targets (Fig. S1E,F), support the notion of common PP2A-related mechanistic basis of regulation.

Together, these data demonstrate that PP2A regulates a very large fraction of cancer cell phosphoproteome. The data also systematically validate the function of CIP2A, PME-1, and SET as cellular inhibitors of serine/threonine dephosphorylation, and provide the first systemic estimate of the degree of their biological redundancy.

### PP2A-regulated cellular processes

We used g:Profiler software ^41^ to systematically analyze the most enriched processes in the differentially regulated phosphoproteomes of PPP2R1A, CIP2A, PME-1, and SET. The identified PP2A target processes were found to cover large spectrum of cellular functions (Fig. 2A), but to contain large number of proteins for which there has not previously been evidence for PP2A-mediated regulation (Table S1).

**Fig 2.**
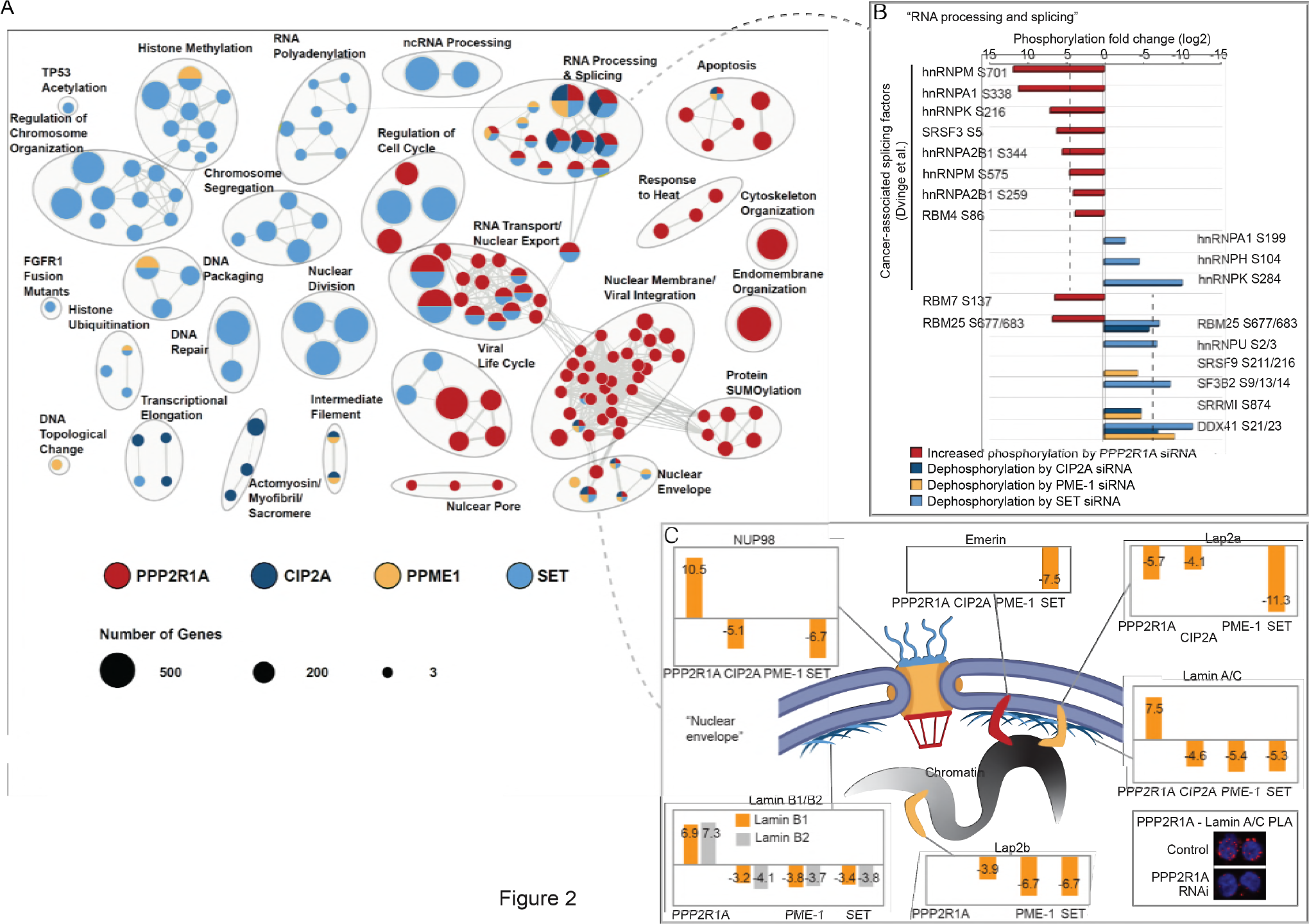
Pathway analysis of PP2A dephosphorylome. A) Pathway enrichment analysis of the differentially regulated phosphopeptides for each condition. B) Differentially regulated phosphosites on proteins in the enriched category “RNA Processing and splicing”. PP2A regulation by PPP2R1A and SET impact phosphorylation of a number of cancer-associated splicing factors. C) Differentially regulated phosphosites on proteins in the enriched category “Nuclear envelope organization” indicates coordinated regulation of nuclear lamina function by PP2A. Proximity ligation assay between PPP2R1A and Lamin A/C is shown in the insert.

Among targets involved in “regulation of cell cycle” we identified examples indicating cross talk between human tumor suppressors PP2A and TP53. SET depletion inhibited phosphorylation of TP53 (serine (S)392; Table S1) and both PPP2R1A and SET regulated MYBBP1A (S1303, S1308), which is involved in TP53 activation via acetylation and functions as a tumor suppressor for RAS-induced transformation ^42^. Another SET target PML (S518, S527) instead controls TP53 acetylation and transcriptional activity ^43^. Other novel SET-specific targets in cell cycle and mitosis regulation included CDC23 (S588, S596), anaphase-promoting complex (APC) subunit CDC26 (S42), and mitotic aurora kinase target BOD1L (S482, S2779, S2964, S2986).

Among the splicing factors with established functional relevance in cancer ^44^, we identified a total of 6 proteins and 8 phospho-sites that were significantly regulated by PPP2R1A (Fig. 2B, and Table S1). While these cancer-relevant splicing factors were predominantly regulated by PPP2R1A, a distinct set of mRNA processing and splicing factors was found to be dephosphorylated upon depletion of the PIPs (Fig. 2B).

Among “nuclear envelope” proteins, we confirmed the regulation of many established LMNA PP2A target sites (Table S1 and Fig. 2C). This is consistent with the associations of CIP2A ^31^, PME-1 ^45^, and PPP2R1A (Fig. 2C insert) with LMNA. We also discover phosphoregulation of LMNB1, LMNB2 and nuclear pore proteins (NUPs), including NUP98 by PPP2R1A and PIPs (Fig. 2C). Additionally, SET depletion induced the dephosphorylation of LAP2A, LAP2B and EMERIN, whereas CIP2A and PME-1 exhibited more selective roles in the phosphoregulation of these nuclear lamina anchor proteins (Table S1 and Fig. 2C).

In summary, our analysis uncovers hundreds of novel PP2A targets associated with large number of cellular processes, many of which are deregulated in cancer. Based on this data, PP2A’s role in controlling cell behavior extends far beyond its currently understood role in controlling kinase pathways, mitosis ^46^, and critical signaling nodes in cancer.

### Chromatin and MYC-associated PP2A targets

SET directly binds to catalytic PP2Ac subunit of PP2A complex, and inhibits the phosphatase activity against various peptide substrates ^24,25^. SET is also a member of inhibitor of acetyltransferases–complex (INHAT) that binds histones, thus directly preventing their acetylation ^47^. In addition to binding core histones, SET has been reported to function as linker histone chaperone ^48^. Interestingly, in our data, functions such as “chromosome organization”, “chromatin modifications”, and “DNA methylation” (Fig. 2A) were strongly enriched in the SET targets. As an example, we identified a number of epigenetic regulators to be significantly dephosphorylated in SET-depleted cells: SMARCC2 (S283), KDM1A (S166, S849), SETD2 (S2080, S2082), HDAC1 (S393, S421, S423), CHD3 (S1601, S1605), MTA1 (386, 576) DNMT1 (S714), DOT1L (S1001, S1009) and BRD1 (S1052, S1055) (Fig. 3A, Table S1). In addition to SET, proteins in the Nurd complex were also found to be regulated by PPP2R1A and PME-1 depletion (Fig. 3A). These results indicate that inhibition of PP2A-mediated protein dephosphorylation may play a more important role in SET’s chromatin-associated functions than previously anticipated.

**Fig 3.**
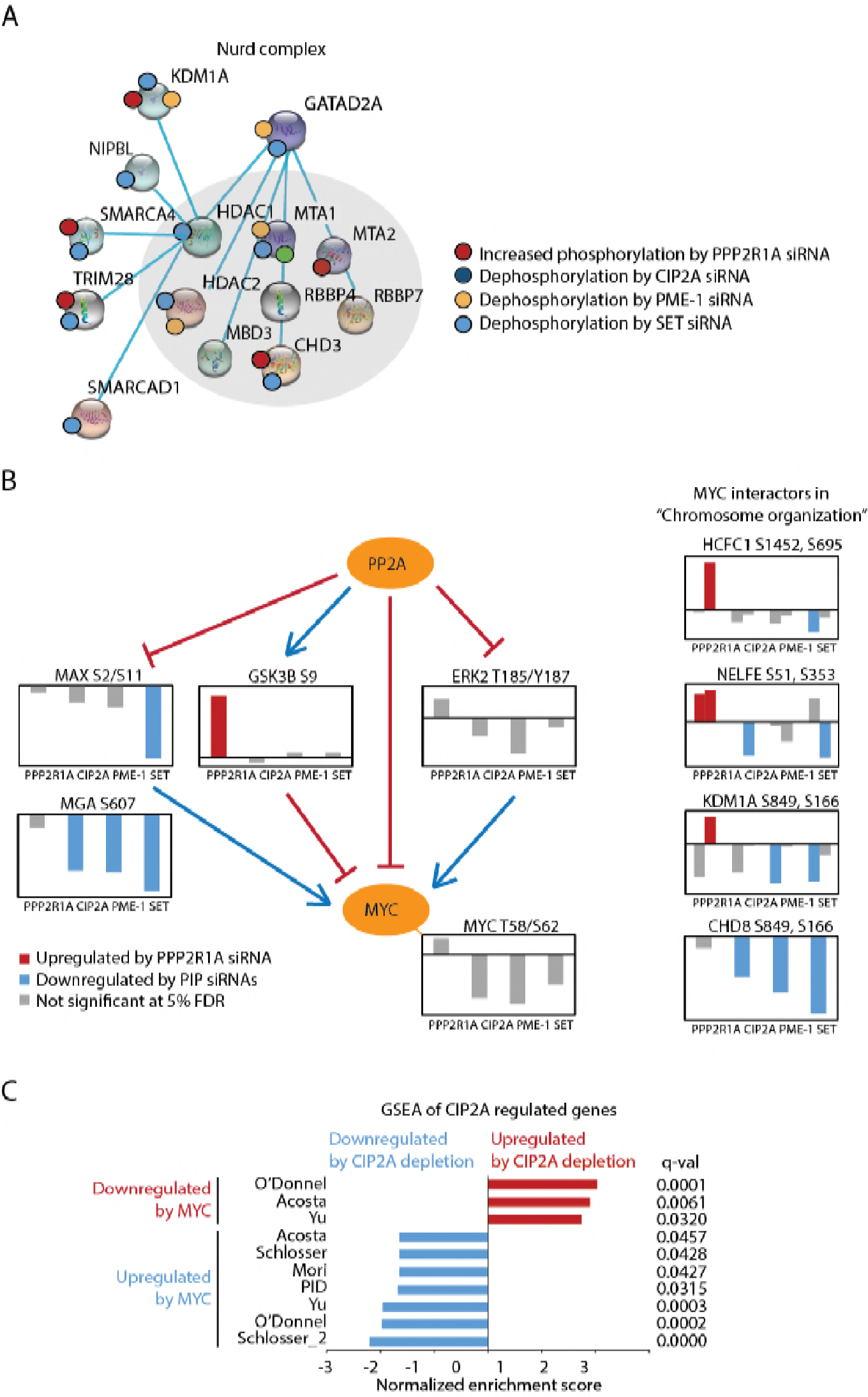
Chromatin, and MYC-associated PP2A targets. A) Nurd complex as an example of a chromatin-associated protein complex regulated by SET and PPP2R1A. B) Graphical representation of PP2A-controlled mechanisms associated with regulation of MYC protein stability and transcriptional activity. C) A summary of impact of CIP2A depletion on MYC-associated gene expression programs based on GSEA. The individual GSEA results are shown in figure S2B.

The best understood chromatin-associated target for PP2A is transcription factor MYC, and regulation of MYC activity by PP2A has primarily been attributed to dephosphorylation of serine 62 ^9,11^. Our data confirmed the approximately 4-fold reduction in the phosphorylation of the peptides containing both T58 and S62 via the depletion of CIP2A, PME-1, or SET (Fig. 3B; Table S1). On the other hand, depletion of PPP2R1A induced GSK3B (S9) phosphorylation (Fig. 3B). This phosphorylation inhibits GSK3B-mediated MYC T58 phosphorylation and thus result in MYC stabilization ^49^. We also confirmed proteasome-mediated MYC degradation due to CIP2A depletion (Fig. S2A).

The pivotal impact of PP2A on the transcriptional activity of MYC was confirmed from the gene set enrichment analysis (GSEA) of the differentially regulated transcripts in CIP2A-depleted HeLa cells (Fig. 3C, S2B). In this regard, we also identified novel potential mechanisms for direct regulation of the transcriptional activity of MYC. SET was found to regulate phosphorylation of MYC dimerization partner MAX at S2/S11 (Fig. 3B; Table S1). The phosphorylation of these sites determines the kinetics of MYC/MAX dimer DNA binding. Furthermore, depletion of CIP2A, PME-1, or SET induced dephosphorylation of the MAX gene-associated protein MGA (S607)(Fig. 3B). MGA is a functional antagonist of MYC, and a recent study demonstrated that MGA loss-of-function mutations and focal MYC amplification are mutually exclusive in human lung cancer ^50^. Interestingly, consistently with newly characterized role of PP2A in regulating chromatin remodeling factors (Fig. 3A), PP2A modulation also influenced the phosphorylation of high-confidence MYC-interacting partners, including CHD8, KDM1A, RDBP, and HCFC1 (Fig. 3B; Table S1); which all display coincident chromatin binding with MYC^51^. Lastly, among the SET-regulated spliceosome targets (Fig. 2B), SF3B2 is a homologue of SF3B1 that is essential for the survival of cancer cells with high MYC activity ^52^, displaying yet another potential level at which PP2A and MYC activities may converge.

These results reveal SET as a chromatin-associated PP2A inhibitor protein. Furthermore, the newly identified regulation of MYC DNA binding mechanisms (MAX and MGA), and of chromatin modifying MYC interactors, provide novel mechanistic insight into how PP2A modulates MYC activity in cancer.

### Unidirectionality paradigm of phosphoregulation by PP2A

It is generally assumed that phosphorylation sites are under constant equilibrium between kinase and phosphatase activities, and can thus be bidirectionally modulated towards either increased or decreased phosphorylation. However, we made an intriguing observation that the majority of PPP2R1A-regulated peptides (211/320 targets, 66%) were not significantly regulated by the inhibition of CIP2A, PME-1, or SET (Fig. 4A). The non-overlapping targets of PPP2R1A and PIPs were reflected also at the level of PP2A-regulated processes that were mostly dominated either by PPP2R1A or by one of the PIPs; and most dominantly by SET (Fig. 2A). Even among the processes such as “RNA Processing and Splicing” that were globally controlled by both PPP2R1A and PIPs (Fig. 2A), most of the individual peptides were significantly regulated only by either PPP2R1A or by PIPs (Fig. 2B).

**Fig.4.**
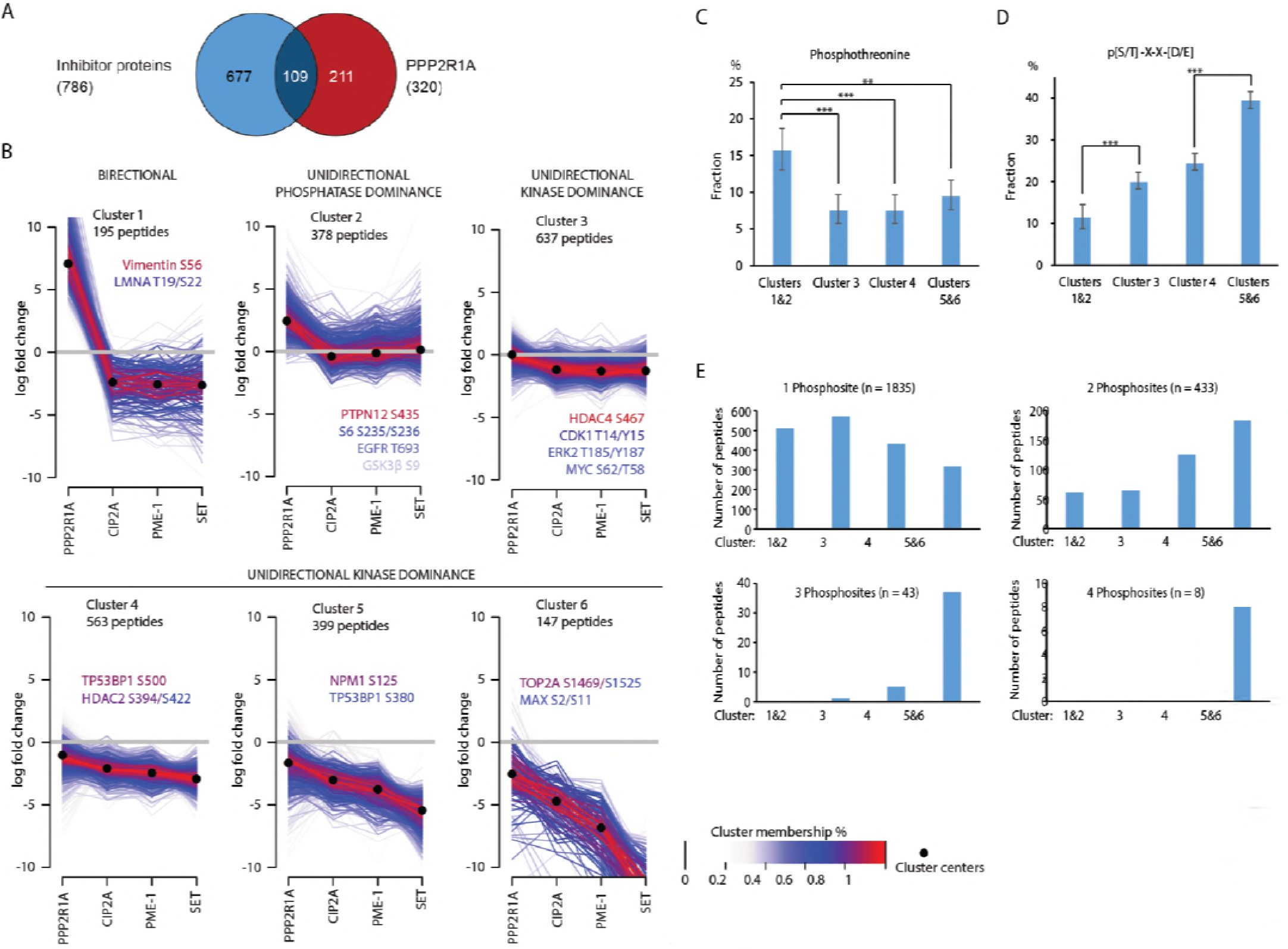
Soft clustering reveals undirectionality of PP2A dephosphorylome regulation. A) Venn diagram presentation of a degree of overlap between PPP2R1A and PIP-regulated peptides at 5% FDR. B) Soft clustering analysis of the high confidence data set into 6 clusters. Cluster membership percentage is indicated by a color scale and representative peptides are listed inside the plots. Cluster centers (indicated by black dots) exhibit mainly unidirectional regulation, upregulation in cluster 1 and downregulation in clusters 3-6. Representative examples of target peptides in each cluster are indicated inside the panels, and colour coded in relation to centrality in the cluster behaviour. C) Fraction of phosphothreonines of all phosphorylation sites in each cluster. D) Fraction of acidophilic kinase targets in each cluster. C,D) Error bars represent 95% confidence intervals. ** represents p<0.01 and *** p<0.001 in Chi2 test. E) Number of peptides with 1, 2, 3, or 4 phosphosites in each cluster.

To further examine these provoking results, and to exclude the possibility that these observations were caused by the selection of the statistical criteria for the differentially regulated peptides presented in figures 1 and 2, we performed a soft clustering analysis ^53^ of the entire dataset of consistently detected peptides. Consistently with the notion of non-overlapping targets of PPP2R1A and PIPs, in five of the six clusters (clusters 2-6; Fig. 4B), the majority of the peptides were subject only to unidirectional regulation toward either increased phosphorylation by PPP2R1A inhibition (cluster 2), or increased dephosphorylation by inhibition of PIPs (clusters 3-6; Fig. 4B). The term unidirectionality is used hereafter to describe a situation where a peptide phosphorylation, or cellular function, is subject to regulation to only one direction by PP2A modulation. On the other hand, cluster 1 was the only cluster in which the same peptides exhibited increased phosphorylation upon PP2A inhibition, and dephosphorylation upon inhibitor protein depletion (i.e., bidirectionality). These results strongly indicate that a large fraction of PP2A target phospho-sites exist in either a nearly fully phosphorylated (kinase dominance; clusters 3-6) or dephosphorylated (phosphatase dominance; cluster 2) state. Although this unidirectional pattern of phosphorylation regulation has not been previously addressed in relation to phosphatase biology, it is consistent with the reportedly bimodal distribution of global phosphorylation site occupancy ^54,55^.

The unidirectionality paradigm of PP2A target regulation is further supported by analysis of the data for factors with previously reported association with high - or low phosphate occupancy. Based on previous work, a higher PP2A/kinase activity ratio is associated rather with threonines than serines ^56–58^, and acidophilic kinase target sites exhibit high phosphate occupancy ^55^. Additionally, phosphorylations on sites in close proximity to each other tend to be phosphorylated in a coordinated fashion by the same kinase, which suggests kinase dominance in regions with high occupancy of adjacent phosphosites ^59^. To analyze our data according to these parameters, clusters 1 and 2 (cluster 1&2 hereon), and clusters 5 and 6 (cluster 5&6 hereon) were combined to yield 4 nearly equally sized clusters. By using these four clusters, we found that phosphothreonines were indeed enriched in phosphatase-dominant peptides (Fig. 4C), whereas acidophilic kinase targets (Fig. 4D), and clustered phosphorylation (peptides with 3 or 4 phosphates) (Fig. 4E) were enriched in kinase-dominant peptides (clusters 4, 5&6). These results are fully in concert with the unidirectionality being a dominant feature in PP2A target regulation.

Collectively, these analysis reveal unidirectionality paradigm of PP2A target regulation. These novel insights are not only important for general understanding of rules of PP2A-mediated phosphoregulation, but also for realization that all phosphoregulation in cancer cells is not equally susceptible to modulation of PP2A activity. Rather, phosphatase-dominant targets would primarily respond to pharmacological PP2A inhibition ^38,60^, whereas various PP2A reactivation therapies ^14,15^, would primarily affect kinase-dominant targets.

### Unidirectional control of canonical pathways

To address how the unidirectionality observed at the peptide level translates to control of cellular activities, we analysed the enrichment of cellular processes and pathways between PP2A target clusters. At the level of cellular process, most of the processes did associate with certain cluster, and followed the pattern of unidirectional regulation (Fig. S3). However, the processes identified at the peptide level to be regulated by both PP2A inhibition and activation, such as RNA splicing, displayed bidirectional regulation also when assessed at the cluster level (Fig. S3). This analysis clearly indicate that the directionality concept of PP2A target regulation is not restricted to individual peptide level, but is relevant to regulation of different cellular processes.

Enrichment of different signalling pathways and kinase targets between different clusters was evaluated by Ingenuity pathway analysis and NetworKIN kinase target prediction tool. As was case with cellular processes, also different signalling pathways displayed notably different distribution between the clusters. Enrichment of PTEN signaling pathway and AKT targets in cluster 1&2 (Fig. 5A, B), suggests that they are mostly under constitutive dephosphorylation by PP2A. Similarly, Phospholipase C (PLC) signaling, and specifically PKC targets, seem also to follow phosphatase-dominated control as they were found to associate with clusters 1-3 (Fig. 5A, B). Interestingly, SET phosphorylation by PKC promotes PP2A inhibition ^61^, which implies the presence of a positive feedforward loop that promotes PKC signal propagation. In contrast, ERK/MAPK signaling pathway components were enriched in phosphatase-dominant cluster 3 (Fig. 5A, B), indicating a clear difference in directionality rules by which different PP2A target pathways are controlled.

**Fig 5.**
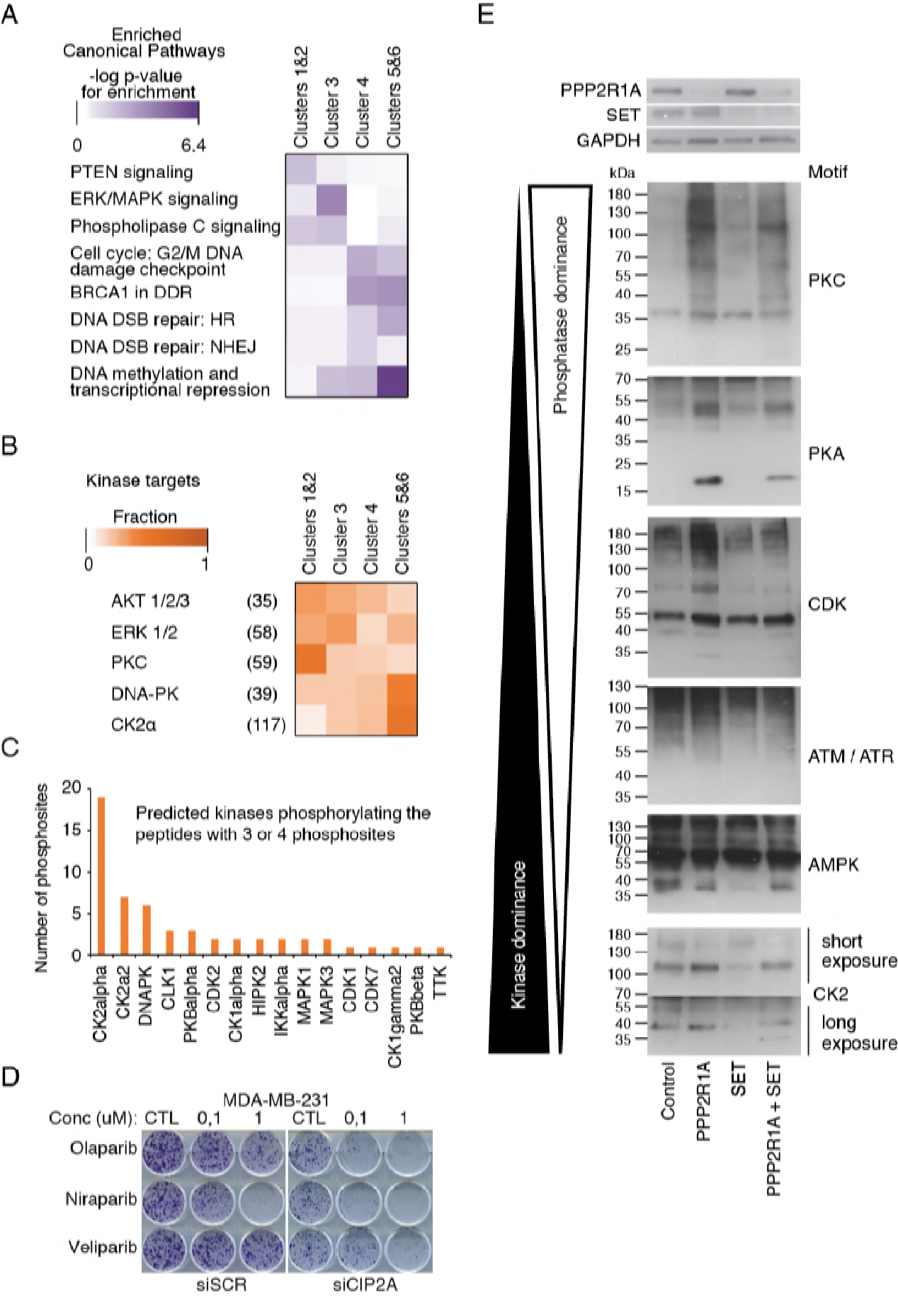
PP2A controls DNA repair mechanisms and PARP inhibitor resistance. A) Enrichment of selected canonical pathways into different clusters. B) Enrichment of selected kinases’ targets into different clusters. C) Prediction of kinases phosphorylating PP2A dephsophorylome peptides with 3 or 4 phosphosites. D) Response of CIP2A-depleted and control MDA-MB-231 cells to indicated PARP inhibitors. E) Analysis of phosphorylation of targets of different kinases by phosphomotif antibodies arranged from the most PPP2R1A responsive (phosphatase dominance) to most SET responsive (kinase dominance) phosphoproteins in descending order.

The kinase target motif analyses also revealed that PP2A-regulated CK2 target sites are enriched on clusters 5&6, that constitute of peptides regulated unidirectionally by PIPs, and dominated by SET regulation (Fig. 5B). This is consistent with previous data indicating functional antagonism between PP2A and CK2 function ^62^, and with role of CK2α as the predominant acidophilic kinase phosphorylating the kinase dominant peptides with multiple phosphorylations (Fig. 5C).

The dephosphorylation-based regulation of DNA repair mechanisms was strongly apparent from the analysis. Canonical DNA repair-associated pathways (Fig. 5A), and DNA-PK targets (Fig. 5B) were enriched in clusters 4-6; and DNA-PK was among the most enriched kinases predicted to phosphorylate kinase dominant sites with multiple phosphorylations (Fig. 5C). Consistent with high basal level of phosphorylation, these clusters consist of proteins that are dephosphorylated upon PIP depletion, but whose phosphorylation cannot be further increased by PPP2R1A inhibition. As all three enriched DNA repair pathways are implicated in PARP inhibitor (PARPi) sensitivity ^63^, we analyzed PARPi sensitivity in CIP2A-depleted MDA-MB-231 triple-negative breast cancer (TNBC) cells. Notably, CIP2A depletion resulted also in a robust “BRCAness” fenotype demonstrated by sensitization to three different PARPi drugs at 0.1 μM concentration (Fig. 5D). These data are fully consistent with recently emerging data indicating that DNA damage response is a cellular process in which protein dephosphorylation plays a particularly important regulatory role ^64^.

The unidirectionality rules revealed by the MS results were confirmed by phospho-motif antibodies. Consistently with the directionality principle, PPP2R1A depletion increased the phosphorylation of different phospho-motifs than those dephosphorylated by SET depletion (Fig. 5E). Moreover, double knockdown of PPP2R1A and SET rescued majority of dephosphorylations by SET depletion, but had less impact on increased phosphorylation by PPP2R1A depletion (Fig. 5E), which is consistent with SET modulating PP2A function, and not vice versa.

In summary, these data strongly indicate that unidirectionality of phosphoregulation by PP2A affects entire pathways and biological processes, and thus it constitutes a biologically relevant signaling principle. As exemplified by identification PP2A inhibition as PARPi resistance mechanism, this novel understanding may help in identification of both target processes, as well as clinically relevant therapy combinations for emerging PP2A-targeted cancer therapies.

### PP2A activity follows the intracellular cytoplasm-nuclear gradient

In addition to directionality paradigm, cluster analysis of dephosphorylome data revealed several indications for intracellular cytoplasm-nuclear gradient of PP2A target regulation. To examine this previously uncharacterized aspect of PP2A biology, we assessed subcellular protein localization information of the target proteins in different clusters. Notably, the analysis revealed a significant difference in subcellular localization between the targets in different clusters, where the targets from cluster 1&2 (i.e., the phosphatase-dominant peptides) were dominantly cytoplasmic, whereas the kinase-dominant targets (clusters 4-6) were highly enriched in nuclear proteins (Fig. 6A). Specifically, the regulation observed in clusters 4 and 5&6, in which PME-1 and SET exhibited a stronger influence on phosphorylation than CIP2A (Fig. 4A), was consistent with the subcellular localization patterns of these PIPs (Fig. 6B). The correlation between localization of CIP2A, PME-1, or SET, and the targets they regulate, was apparent also at the level of the differentially regulated peptides, whereas the targets significantly regulated by PPP2R1A depletion were equally distributed between cytoplasm and nucleus (Fig. 6C). As additional examples, the enrichment of PKC targets in the clusters 1&2, and CK2α targets in the cluster 5&6 (Fig. 4B, 6D), is consistent with the previously reported predominantly cytoplasmic and nuclear localization of their targets, respectively ^65^. Further, nuclear pathways involved in cell cycle checkpoints and, DNA repair and transcriptional repression were associated with clusters 4-6 (Fig. 4A), that are most dominantly regulated by the nuclear PIPs PME-1 and SET (Fig. 4A, 6A).

**Fig 6.**
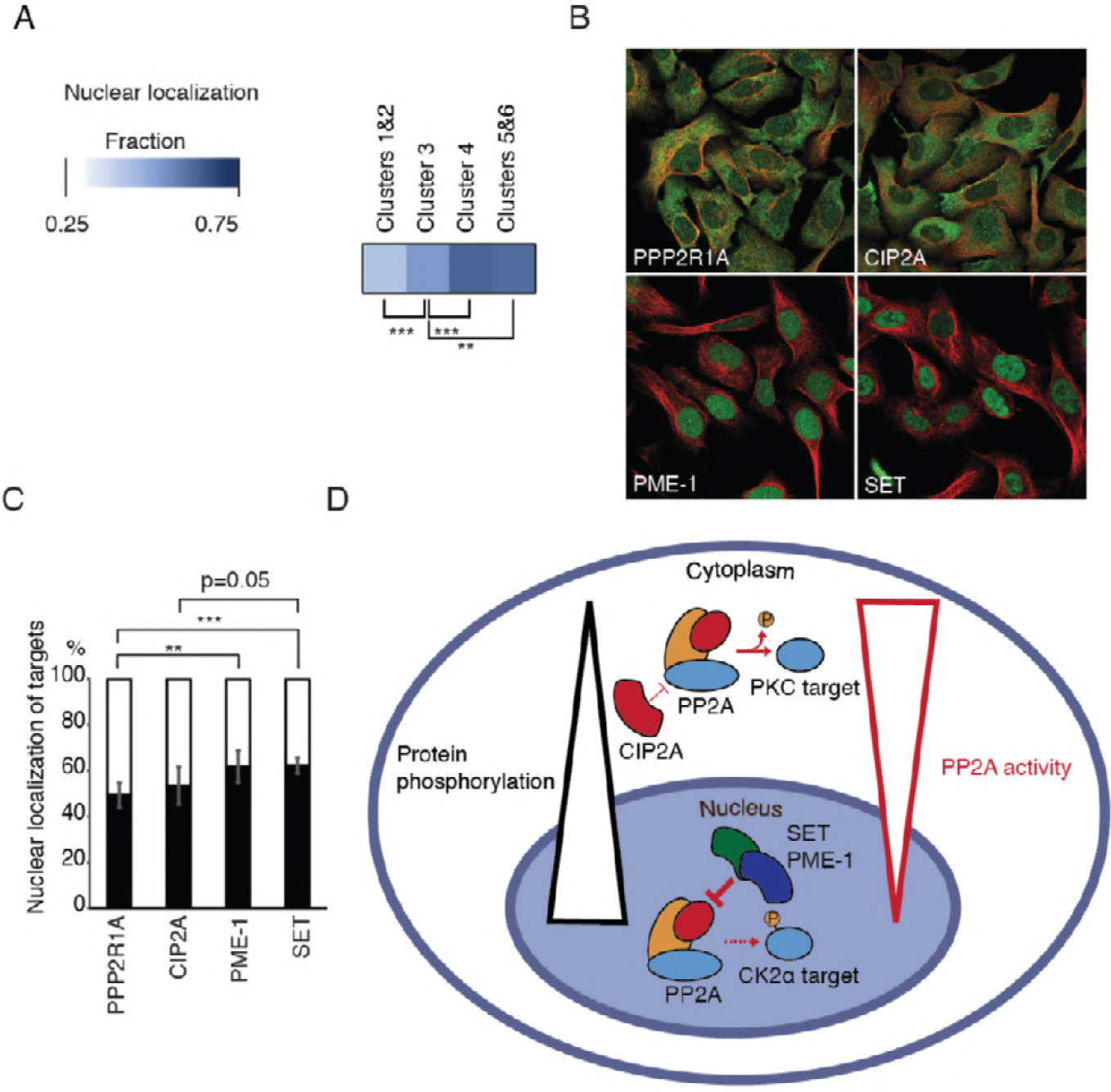
PP2A activity follows the intracellular cytoplasm-nuclear gradient. A) Analysis of fraction of nuclear proteins in each cluster demonstrates association between nuclear localization and unidirectional kinase dominance (Clusters 4-6). B) Subcellular localizations of PPP2R1A and PIPs, retrieved from www.proteinatlas.org. PPP2R1A: CAB018599: U-2 OS, image 2, CIP2A: HPA039570: U-2 OS, image 1, PME-1: CAB004541: U-2 OS, image 1, SET: HPA063683: U-2 OS, image 1) C) Fraction of nuclear proteins in the proteins with differentially regulated phosphopeptides for each condition. In panels A and C, ** denotes p<0.01 and *** denotes p<0.001 for chi^2^-test. D) Schematic presentation of intracellular cytoplasm-nuclear gradient of PP2A activity. Based on low association between nuclear proteins and PP2A dephosphrylome clusters 1&2 it is apparent that cytoplasmic proteins are under higher dephosphorylation activity than nuclear proteins that associate with clusters 4-6 with tight kinase dominance.

Together these data strongly indicate that PP2A activity is not uniformly distributed in the cell, but comprises of a cytoplasm-nuclear gradient in which PP2A inhibition dominates in the nucleus, whereas cytoplasmic balance is shifted towards phosphatase dominance (Fig. 6D). This observed phosphorylation gradient is the opposite of the expected gradient for signals originating at the cell membrane and translocating to nucleus under the assumption of uniform phosphatase activity ^66^. Therefore, the results suggest that efficient signal propagation in cancer cells requires tight inhibition of nuclear PP2A activity by nuclear PP2A inhibitor protein SET and PME-1 (Fig. 6D). Further, these results indicate an important role of the spatial distribution of oncogenic PIPs in signal propagation.

### The PP2A dephosphorylome both quantitatively and qualitatively defines cancer drug responses

Finally, in order to test functional relevance of PP2A dephosphorylome rules characterized in this study, we exposed the PP2A-modulated cells to drug sensitivity and resistance testing (DSRT) platform ^67^. The platform consists of 306 drugs covering all major drug classes from specific kinase inhibitors to DNA-damaging agents and proteosomal inhibitors and thus covers comprehensively different drug-induced biological responses. The cell viability readout of DSRT platform served as a surrogate marker for impact of PP2A modulation to interference with different cellular pathways. The PP2A RNAi treatments were identical to those applied for dephosphorylome profiling and differential drug sensitivity score (ΔDSS) ^68^ was calculated between the treated and control samples. Among 306 tested drugs, 68 were excluded because they elicited no response in HeLa cells, regardless of the siRNA treatments (Table S2).

The shift in the dephosphorylation balance (Fig. 1B, C) was used as an indicator of PP2A dephosphorylome activity, and the drugs were ranked by uncentered Pearson’s correlations between dephosphorylome activity and the changes in DSSs (Fig. 7A). The drugs that exhibited synergy with high dephosphorylome activity appear at the top of “correlation rank” (red), whereas the drugs at the bottom exhibited synergy with PP2A inhibition (blue). Furthermore, the enrichment scores were calculated for selected drug groups in the ranked list (Fig. 7A). The individual drug responses are listed in table S2.

**Fig 7.**
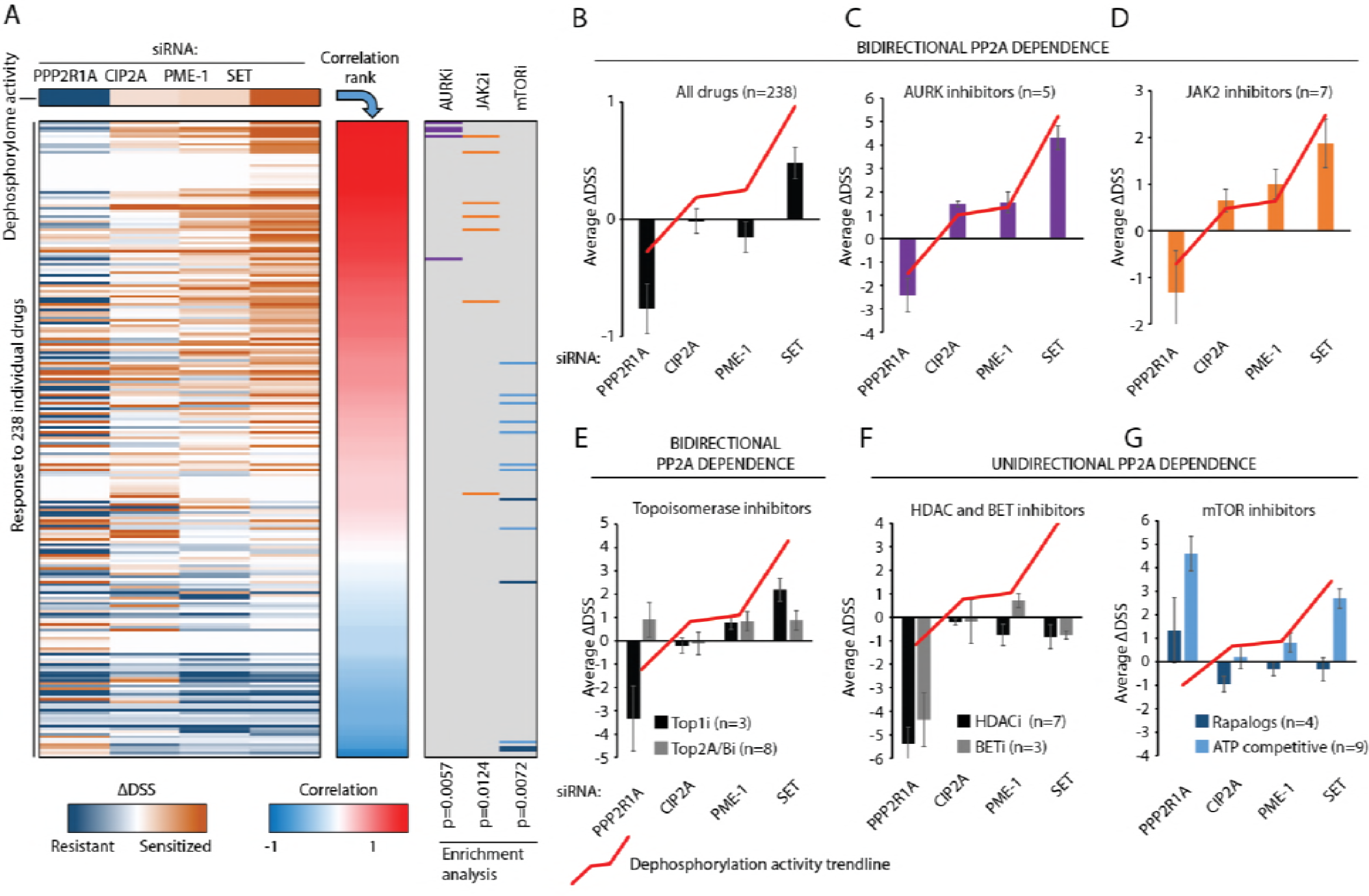
PP2A dephosphorylome activity defines sensitivities across drugs representing diverse modes of action. A) Drug sensitivity scores (ΔDSS) of 238 cancer drugs in HeLa cells are ranked by correlation to PP2A activity index derived from differentially regulated peptides for each gene; represented as the top color bar in A) panel and red line in the subsequent panels. Enrichment of selected drug groups is shown on the right panel B) Average ΔDSS over all 238 drugs, C) Aurora kinase inhibitors, D) JAK2 inhibitors, E) topoisomerase inhibitors, F) HDAC and BET inhibitors, G) and ATP-competitive and rapalog-mTOR inhibitors.

Remarkably, PP2A inhibition by PPP2R1A depletion resulted in increased average drug resistance across all 238 drugs, whereas robust increases in protein dephosphorylation due to the depletion of SET caused the cells to become more drug sensitive on average (Fig. 7A, B). Interestingly, although the depletion of CIP2A and PME-1 did not influence average drug sensitivity (Fig. 7B), their effect on sensitivity to individual drugs was in many cases similar to that of SET, albeit of lower magnitudes (Fig. 7A). This is consistent with the weaker influence of CIP2A and PME-1 on dephosphorylome activity compared to SET (Fig. 1C). The correlation between overall dephosphorylome activity and drug response was most evident with Aurora kinase (AURK) inhibitors (p=0.0057 for enrichment), and JAK2 inhibitors (p=0.0142) (Fig. 7A,C, and D). Interestingly, reflecting the complicated crosstalk between PP2A and mitotic kinases where spatial control contributes to dephosphorylation balance, the bidirectional PP2A activity dependency of the AURK inhibitors was not shared by the other antimitotic drugs (Fig. S4A). TOP1 inhibitors, but not TOP2B inhibitors, were identified as another class of cancer drugs whose effect correlated with dephosphorylome activity (Fig. 7E). Further, consistent with other results, PARP inhibitors Olaparib and Rucaparib exhibited clear PP2A-dependency (Table S2).

The unidirectionality of PP2A-mediated regulation of drug sensitivity was evident for HDAC and BET bromodomain inhibitors. While PPP2R1A depletion induces significant resistance across all the HDAC and BET inhibitors, increased dephosphorylation activity due to PIP depletion did not confer any drug sensitization (Fig.7F). This resistance to HDAC and BET inhibition could reflect the apparent role of PP2A in chromatin regulation (Fig. 2A and 3A). The BET inhibitor results are consistent with recently reported role of PP2A inhibition in JQ1 resistance in TNBC ^69^. Another interesting unidirectional PP2A dependency was observed for mTOR inhibitors, which in contrast to the majority of drugs, exhibited synergy with PPP2R1A inhibition (Fig. 7G). The observed sensitization to mTOR inhibitors may reflect an acquired dependency on increased mTOR activity in PPP2R1A-depleted cells, consistent with upregulation of PI3K/AKT/mTOR signaling in these cells (Fig S4B,C). Furthermore, the role of SET in mTOR inhibitor resistance was particularly interesting because only ATP-competitive mTOR inhibitors exhibited sensitization upon SET depletion (Fig. 7G). This finding may be related to the unexpected observation that also SET inhibition caused AKT activation, albeit with differences with PPP2R1A in downstream target phosphorylation (Fig. S4B,C).

Together, these data demonstrate a notable overall correlation between PP2A dephosphorylome activity and cellular response to drugs representing all major drug classes. The most important notions however might be that different drug classes can be classified based on their PP2A dependency of response, and that directionality rules apply also to PP2A-dependent drug responses. These aspect may become relevant for developing combination strategies with the emerging PP2A-targeted therapies aiming either to re-activate ^14,15^, or inhibit ^38,60^ PP2A.

## Discussion

Despite of essential importance of PP2A activity for normal growth and development, as well as its involvement in various diseases, our understanding of PP2A-regulated signaling remains superficial. In this study, we present the largest systematic analysis of PP2A dephosphorylome regulation by the structural PP2A scaffold PPP2R1A, and the three PP2A inhibitor proteins. Overall the study confirmed the widespread impact of PP2A in cellular phosphopregulation, but also revealed unprecedented rules by which PP2A governs cellular serine/threonine phosphorylation. The results also reveal PP2A activity as a general determinant of cellular drug responses across all major drug classes. The insights provided by these results are important for holistic understanding of cell signaling, but also may have translational relevance in light of emerging pharmacological approaches for PP2A activity modulation in human diseases.

Related to specific targets and processes regulated by PP2A, we characterized a number of signaling nodes where PP2A regulates number of proteins that also physically interact with each other. This indicates for a finely-tuned and co-ordinate regulation of function of these complexes by PP2A activity. In addition to a number of directly cancer relevant targets and processes, we also identified many cellular processes that are relevant for all types of cells but for which there thus far has been very little information about role of PP2A. Examples of such processes are mRNA splicing, regulation of nuclear lamina, and chromatin remodelling. Especially interesting is the evidence that inhibition of protein dephosphorylation is an integral part of chromatin-associated functions of nuclear PP2A inhibitor protein SET. Together with previous data indicating strong association of SET with histones, we speculate that this may generate spatially restricted inhibition of dephosphorylation of chromatin remodeling complexes and that SET modulation by emerging small molecule approaches ^70^, may provide novel opportunities for targeting epigenetic gene regulation. Another interesting example of PP2A-regulated processes was “Viral life cycle” (Fig. 2A) which included several viral replication factors and proteins implicated in viral genome transcription (Table S1). PP2A-mediated regulation of viral host proteins is particularly interesting as several viruses encode for PP2A inhibitory proteins that specifically interfere with PP2A functions to promote malignant transformation ^71^.

Directly related to cancer, but also to many physiological processes, is PP2A-mediated regulation of MYC activity. Thus far PP2A’role in MYC regulation has been tightly linked to regulation of MYC stability via GSK3B and phosphorylation of N-terminal degron containing threonine 58 and serine 62 ^11^. We confirm those PP2A-mediated effects, but in addition to that, our results expose an entirely new level of regulation of MYC activity by PP2A-mediated dephosphorylation of MAX, MGA and a number of epigenetic factors that physically associate with MYC at MYC binding sites in chromatin ^51^. These results perfectly explain why, despite of its rather modest impact on MYC protein levels (Fig. S2A), CIP2A depletion has such profound effect on MYC-mediated transcriptional activity both *in vitro* (Fig. 3C), and *in vivo* ^31^. As inhibition of CIP2A blocks MYC’s proliferation promoting activity *in vivo* ^31^, but PP2A reactivation, either by CIP2A depletion or pharmacologically, do not cause detrimental physiological effects ^10,14,29,31^, these results highlight the rational of targeting MYC hyperactivity via PP2A modulation. Notably, although CIP2A is mostly a cytoplasmic protein, the profound effects of CIP2A on MYC activity can be explained by previously published evidence for spatially restricted nuclear lamina-associated domains (LADs) in which CIP2A regulates MYC phosphorylation ^31^. With our new data indicating widespread effects of PP2A also on nuclear lamina, these data present another elegant example of coordinated impact of PP2A in cell biology.

In addition to novel information of PP2A-regulated processes and targets, the data also reveal important general concepts related to phosphosignaling. For example, we demonstrate previously unknown differential spatial distribution of kinase-and phosphatase-dominant targets (Fig. 6A, D) and the influence of the amino acid context on the propensity for dephosphorylation (Fig. 4C-E). The data all together indicate that cellular responses to specific PP2A manipulations are determined by whether the certain phosphosites are either in a kinase-or phosphatase-dominant state. Importantly, we further demonstrate that these rules do not only apply to individual phosphopeptides, but that entire cellular processes are under either kinase or phosphatase dominance (Fig. 5A,B, S3). In addition to broad conceptual importance of appreciating the directionality rules of PP2A-mediated phosphoregulation, this novel information is essential in regards to understanding that not all targets and cellular process are equally amendable for modulation by emerging PP2A-targeted therapies.

Previously PP2A activity has been shown to modulate cellular responses to certain individual drugs. However, the extent of importance of PP2A activity for global drug response observed in this study is unprecedented as compared to any other signaling mechanism studied as yet. Among the clearest examples of PP2A-dependent drug classes were AURK and JAK2 kinase inhibitors, and epigenetic HDAC and BET inhibitors. PP2A activity correlation with AURK inhibitor response is consistent with a critical role for PP2A in regulation mitotic phosphorylation events ^46^. Whereas higher PP2A activity correlated in average with better drug sensitivity across all major drug classes, also opposite patterns where PP2A inhibition sensitized to drug effects were observed. Together these findings clearly highlight the further need to understand the PP2A-dependency of major drug classes in order to design more effective combination therapy approaches including pharmacological PP2A activity modulators. Consistently with this idea, we have recently demonstrated a potent *in vivo* synergy between pharmacological PP2A reactivation and MEK inhibitors in the preclinical model of mutant KRAS-driven lung cancer ^72^. Notably, the unidirectionality paradigm of PP2A dephosphorylome regulation became very apparent in the drug response data where we observed many examples of both bi-and unidirectional PP2A dependencies among different drug classes. Thereby, our results not only identify individual drug-gene correlations, but potentially reveal globally relevant rules that may help to design better future combination therapy strategies with emerging pharmacological PP2A modulators.

CIP2A, PME-1, and SET constitute a novel class of human oncoproteins ^4,19,20^. Previous work clearly indicated that the oncogenic function of these PIPs is dependent on their PP2A inhibitory function ^10,26–28^. These proteins also constitute very promising novel cancer therapy targets. However, thus far there has not been systematic understanding of role of these proteins in phosphoproteome regulation, or about their biological redundancies. Based on the obtained data, the differential expression that occurs across human cancer types and the differences in subcellular compartmentalization between PIPs may play important roles in determining their biological redundancy. In addition to these qualitative aspects, the quantitative differences in dephosphorylome regulation by CIP2A, PME-1, and SET are likely explainable by the different functional modes through which these proteins modulate PP2A function ^21,23,24^. It is conceivable that the truly non-redundant targets between CIP2A, PME-1, and SET, for the first time systematically analyzed in this study, may explain why only some of these proteins are essential for normal growth and development ^29,32^. These qualitative differences may also be important attributes in assessing the potential and safety of targeting these proteins as a therapeutic approach for the treatment of cancer and other diseases.

In summary, this study provides the largest currently available resource for understanding the target processes associated with the critical human tumor suppressor PP2A. The data reveal unprecedented role of PP2A in various cellular processes and a number of novel connections between PP2A and oncoproteins, as well as with other tumor suppressors. This study also provides the first systematic analysis of the biological redundancy and downstream targets of clinically relevant human oncoproteins CIP2A, PME-1, and SET. All together, these discoveries provide the basis for better understanding of how PP2A functions as a master regulator of cellular Ser/Thr phosphorylation, as a tumor suppressor and as a modulator of cellular drug responses. With emerging PP2A-targeted therapies, this data will also provide important insights into understanding what processes might be most amendable for modulation by PP2A reactivating small molecules and what are the drugs the PP2A modulating drugs should be combined to yield most efficient cellular responses. On a conceptual level, our data provides important novel insight into how serine/threonine phosphatases coordinate cell phosphosignaling and drug responses.

## Materials and methods

### Cell culture and transfection

HeLa cells were culture in DMEM containing 10% FBS, 2 mM glutamine, 50 I.U./ml penicillin, and 50 μg/ml streptomycin. The cells were transfected with Oligofectamine (250 nM siRNA) according to manufacturer’s protocol (Life Technologies). CIP2A was targeted with 4 siRNAs and the other genes with 3 siRNAs. Knockdown was validated by western blotting and RT-qPCR. siRNA sequences are presented in Table S3

### Phosphoproteomics

Sample preparation, LC-MS/MS analysis and the pairwise normalization for label free quantitative phosphoproteomics were performed according to the previously published protocol ^37^. Each siRNA treatment was performed in triplicates for HeLa cells. Samples were collected 72 hours after the transfection. The directionality of regulation by PPP2R1A and PIPs in the HeLa samples was robust to the selection of normalization method (Figure 1B, S1C), however, we observed positive skewing of the fold change distribution by PPP2R1A depletion and negative skewing by depletion of PIPs (Figure 1C, S1C). This is likely a result of upregulation and downregulation, respectively, of significant fraction of phosphosites in these samples (Figure 1C) and supports the selection of pairwise normalization. The directionality of regulation was also consistent across different siRNAs (Figure S1D). The HeLa cell data consists of 2 sets analysed at different times and the peptides displaying batch effect were filtered out. Specifically, linear regression model residuals in phosphopeptide sample triplicates abundances displayed bimodality after imputing smallest non-zero values for non-detected abundances, coupled with log2-transformation, and the phosphopeptides displaying high variance were filtered out before downstream analysis. The differential expression of the filtered dataset was then examined using the popular variance-pooling linear regression R-package limma ^73^, where each case triplicate was compared to the corresponding scrambled control triplicate

### Soft clustering, pathway analyses, motif enrichment

Soft clustering was perfomed in R using Mfuzz package ^74^. The following filtering steps were performed for pathway enrichment analysis: First, a list of unique proteins was constructed by selecting the peptide with the lowest p-value for every protein. Second, unique proteins were filtered using a FDR-corrected p-value cutoff (q ≤ 0.05). Third, the final list of proteins was ranked by the significance (p-value) for subsequent pathway enrichment analysis. Analysis of phosphopeptide clusters was carried out similarly. Clusters were compiled by assigning proteins to clusters based on maximum probability. For proteins with multiple identified peptides, we selected the peptide with highest cluster membership (Table S1) to represent the protein and thus obtained non-redundant lists of unique proteins. The final lists of proteins were ranked according to the cluster membership probabilities. Pathway enrichment analysis was performed using the g:Profiler R package ^41^. We selected biological processes of Gene Ontology and molecular pathways of Reactome as data sources for pathway analysis and filtered the other sources. We used the default method of multiple testing correction and filtered results by corrected significance cut-off of q<0.05. We also limited gene sets to include between 3 and 500 genes. The background list for pathway enrichment analyses included the unique list of all proteins from our dataset combined with all known human phosphoproteins. The phosphoproteins were retrieved from the PhosphoSitePlus database ^75^, and were filtered to include previously published results (“MS_LIT”, “LT_LIT”). Protein names in the background set were converted to gene IDs of the Ensembl database using the g:Convert feature of g:Profiler. This procedure provided a list of enriched pathways for each condition. The lists were merged into a unified pathway-condition table and visualised with the Enrichment Map app ^76^ in Cytoscape ^77^. Finally, we manually reviewed the Enrichment Map and grouped into the most representative functional themes.

Enrichment of canonical pathway components in the clusters, as well as subcellular localization of the proteins, were analysed with Ingenuity Pathway Analysis, spring 2015 release version (QIAGEN). Kinase target predictions were performed with NetworKIN and NetPhorest ^78^. Enriched phosphosite motifs were analysed with motif-x ^79^. Proteins with B56 binding motif ^40^ were retrieved from http://slim.ucd.ie/pp2a/ using default search parameters.

### Drug sensitivity testing

For the high-throughput DSRT-analyses, cells were transfected 3 days prior to plating them on the drug-containing plates at confluency of 1000 cells/well. The subsequent analyses were carried out as described previously ^67^. The differential drug sensitivity score (ΔDSS) ^68^ was calculated by comparing the response of each siRNA-depleted sample to the average of control samples.

For colony formation assays, 3000 cells in each well on 6-well plate were plated one day after transfections together with drugs, and the assays were performed as described previously ^37^. The cells were stained crystal violet similarly to the colony formation assay and the coverage was assessed visually.

### Enrichment of drug groups

The numbers of upregulated phosphopeptides for each gene were substracted from the number of downregulated phosphopeptides and the derived difference values were used as PP2A activity index. Uncentered Pearson’s correlation coefficients were calculated between the PP2A activity index and DSS drug response values for each drug. Drugs were then ranked by correlation with PP2A activity index. Enrichment scores for selected drug groups in the ranked lists were calculated similarly to Gene Set Enrichment Analysis ^80^

### Cell staining

Proximity ligation assay was performed with Duolink kit (Sigma-Aldrich) according to manufacturer’s protocol.

## References

1 Kong, M., Ditsworth, D., Lindsten, T. & Thompson, C. B. Alpha4 is an essential regulator of PP2A phosphatase activity. Mol Cell 36, 51–60, doi:10.1016/j.molcel.2009.09.025 (2009).

2 Apostolidis, S. A. et al. Phosphatase PP2A is requisite for the function of regulatory T cells. Nat Immunol 17, 556–564, doi:10.1038/ni.3390 (2016).

3 Martin, L. et al. Tau protein phosphatases in Alzheimer’s disease: the leading role of PP2A. Ageing Res Rev 12, 39–49, doi:10.1016/j.arr.2012.06.008 (2013).

4 Perrotti, D. & Neviani, P. Protein phosphatase 2A: a target for anticancer therapy. The lancet oncology 14, e229–238, doi:10.1016/S1470-2045(12)70558-2 (2013).

5 Hahn, W. C. et al. Enumeration of the simian virus 40 early region elements necessary for human cell transformation. Mol Cell Biol 22, 2111–2123. (2002).

6 Rangarajan, A., Hong, S. J., Gifford, A. & Weinberg, R. A. Species-and cell type-specific requirements for cellular transformation. Cancer Cell 6, 171–183 (2004).

7 Meeusen, B. & Janssens, V. Tumor suppressive protein phosphatases in human cancer: emerging targets for therapeutic intervention and tumor stratification. Int J Biochem Cell Biol, doi:10.1016/j.biocel.2017.10.002 (2017).

8 Ruediger, R., Ruiz, J. & Walter, G. Human Cancer-Associated Mutations in the A alpha Subunit of Protein Phosphatase 2A Increase Lung Cancer Incidence in A alpha Knock-In and Knockout Mice. Molecular and Cellular Biology 31, 3832–3844, doi:Doi 10.1128/Mcb.05744-11 (2011).

9 Junttila, M. R. et al. CIP2A inhibits PP2A in human malignancies. Cell 130, 51–62, doi:10.1016/j.cell.2007.04.044 (2007).

10 Laine, A. et al. Senescence Sensitivity of Breast Cancer Cells Is Defined by Positive Feedback Loop between CIP2A and E2F1. Cancer discovery 3, 182–197, doi:10.1158/2159-8290.CD-12-0292 (2013).

11 Yeh, E. et al. A signalling pathway controlling c-Myc degradation that impacts oncogenic transformation of human cells. Nat Cell Biol 6, 308–318 (2004).

12 Chen, W., Arroyo, J. D., Timmons, J. C., Possemato, R. & Hahn, W. C. Cancer-associated PP2A Aalpha subunits induce functional haploinsufficiency and tumorigenicity. Cancer Res 65, 8183–8192 (2005).

13 Walter, G. & Ruediger, R. Mouse model for probing tumor suppressor activity of protein phosphatase 2A in diverse signaling pathways. Cell Cycle 11, 451–459, doi:10.4161/cc.11.3.19057 (2012).

14 Sangodkar, J. et al. Activation of tumor suppressor protein PP2A inhibits KRAS-driven tumor growth. J Clin Invest, doi:10.1172/JCI89548 (2017).

15 O’Connor, C. M., Perl, A., Leonard, D., Sangodkar, J. & Narla, G. Therapeutic Targeting of PP2A. Int J Biochem Cell Biol, doi:10.1016/j.biocel.2017.10.008 (2017).

16 Eichhorn, P. J., Creyghton, M. P. & Bernards, R. Protein phosphatase 2A regulatory subunits and cancer. Biochimica et biophysica acta 1795, 1–15 (2009).

17 Naetar, N. et al. PP2A-mediated regulation of Ras signaling in G2 is essential for stable quiescence and normal G1 length. Mol Cell 54, 932–945, doi:10.1016/j.molcel.2014.04.023 (2014).

18 Kauko, O. & Westermarck, J. Non-genomic mechanisms of Protein Phosphatase 2A (PP2A) regulation in cancer. The International Journal of Biochemistry & Cell Biology (2017).

19 Khanna, A., Pimanda, J. E. & Westermarck, J. Cancerous inhibitor of protein phosphatase 2A, an emerging human oncoprotein and a potential cancer therapy target. Cancer Res 73, 6548–6553, doi:10.1158/0008-5472.can-13-1994 (2013).

20 Kaur, A. & Westermarck, J. Regulation of protein phosphatase 2A (PP2A) tumor suppressor function by PME-1. Biochem Soc Trans 44, 1683–1693, doi:10.1042/BST20160161 (2016).

21 Xing, Y. et al. Structural mechanism of demethylation and inactivation of protein phosphatase 2A. Cell 133, 154–163 (2008).

22 Muto, S. et al. Relationship between the structure of SET/TAF-Ibeta/INHAT and its histone chaperone activity. Proc Natl Acad Sci U S A 104, 4285–4290, doi:10.1073/pnas.0603762104 (2007).

23 Wang, J. et al. Oncoprotein CIP2A is stabilized via interaction with tumor suppressor PP2A/B56. EMBO Rep, doi:10.15252/embr.201642788 (2017).

24 Arnaud, L. et al. Mechanism of inhibition of PP2A activity and abnormal hyperphosphorylation of tau by I2(PP2A)/SET. FEBS Lett 585, 2653–2659, doi:10.1016/j.febslet.2011.07.020 (2011).

25 Li, M., Guo, H. & Damuni, Z. Purification and characterization of two potent heat-stable protein inhibitors of protein phosphatase 2A from bovine kidney. Biochemistry 34, 1988–1996 (1995).

26 Niemela, M. et al. CIP2A signature reveals the MYC dependency of CIP2A-regulated phenotypes and its clinical association with breast cancer subtypes. Oncogene 31, 4266–4278, doi:10.1038/onc.2011.599 (2012).

27 Kaur, A. et al. PP2A inhibitor PME-1 drives kinase inhibitor resistance in glioma cells. Cancer Res, doi:10.1158/0008-5472.CAN-16-1134 (2016).

28 Neviani, P. et al. The tumor suppressor PP2A is functionally inactivated in blast crisis CML through the inhibitory activity of the BCR/ABL-regulated SET protein. Cancer Cell 8, 355–368 (2005).

29 Ventelä, S. et al. CIP2A promotes proliferation of spermatogonial progenitor cells and spermatogenesis in mice. PLoS ONE 7, e33209, doi:10.1371/journal.pone.0033209 (2012).

30 Come, C. et al. CIP2A Promotes T-Cell Activation and Immune Response to Listeria monocytogenes Infection. PLoS ONE 11, e0152996, doi:10.1371/journal.pone.0152996 (2016).

31 Myant, K. et al. Serine 62-Phosphorylated MYC Associates with Nuclear Lamins and Its Regulation by CIP2A Is Essential for Regenerative Proliferation. Cell reports 12, 1019–1031, doi:10.1016/j.celrep.2015.07.003 (2015).

32 Ortega-Gutierrez, S., Leung, D., Ficarro, S., Peters, E. C. & Cravatt, B. F. Targeted disruption of the PME-1 gene causes loss of demethylated PP2A and perinatal lethality in mice. PLoS ONE 3, e2486 (2008).

33 Park, H. J. et al. Protein Phosphatase 2A and Its Methylation Modulating Enzymes LCMT-1 and PME-1 Are Dysregulated in Tauopathies of Progressive Supranuclear Palsy and Alzheimer Disease. Journal of neuropathology and experimental neurology 77, 139–148, doi:10.1093/jnen/nlx110 (2018).

34 Trotta, R. et al. The PP2A inhibitor SET regulates natural killer cell IFN-gamma production. J Exp Med 204, 2397–2405, doi:10.1084/jem.20070419 (2007).

35 Trotta, R. et al. The PP2A inhibitor SET regulates granzyme B expression in human natural killer cells. Blood 117, 2378–2384, doi:10.1182/blood-2010-05-285130 (2011).

36 Khanna, A. & Pimanda, J. E. Clinical significance of Cancerous Inhibitor of Protein Phosphatase 2A (CIP2A) in human cancers. Int J Cancer, doi:10.1002/ijc.29431 (2015).

37 Kauko, O. et al. Label-free quantitative phosphoproteomics with novel pairwise abundance normalization reveals synergistic RAS and CIP2A signaling. Scientific reports 5, 13099, doi:10.1038/srep13099 (2015).

38 Lai, D. et al. PP2A inhibition sensitizes cancer stem cells to ABL tyrosine kinase inhibitors in BCR-ABL(+) human leukemia. Science translational medicine 10, doi:10.1126/scitranslmed.aan8735 (2018).

39 Longin, S. et al. Selection of protein phosphatase 2A regulatory subunits is mediated by the C terminus of the catalytic Subunit. J Biol Chem 282, 26971–26980 (2007).

40 Hertz, E. P. et al. A Conserved Motif Provides Binding Specificity to the PP2A-B56Phosphatase. Mol Cell 63,686–695, doi:10.1016/j.molcel.2016.06.024 (2016).

41 Reimand, J., Arak, T. & Vilo, J. in Nucleic Acids Res Vol. 39 W307–315 (2011).

42 Mori, S. et al. Myb-binding protein 1A (MYBBP1A) is essential for early embryonic development, controls cell cycle and mitosis, and acts as a tumor suppressor. PLoS One 7, e39723, doi:10.1371/journal.pone.0039723 (2012).

43 Rokudai, S. et al. MOZ increases p53 acetylation and premature senescence through its complex formation with PML. Proc Natl Acad Sci U S A 110, 3895–3900, doi:10.1073/pnas.1300490110 (2013).

44 Dvinge, H., Kim, E., Abdel-Wahab, O. & Bradley, R. K. RNA splicing factors as oncoproteins and tumour suppressors. Nat Rev Cancer 16, 413–430, doi:10.1038/nrc.2016.51 (2016).

45 Pokharel, Y. R. et al. Relevance Rank Platform (RRP) for Functional Filtering of High Content Protein-Protein Interaction Data. Molecular & Cellular Proteomics 14, 3274–3283, doi:10.1074/mcp.M115.050773 (2015).

46 Gelens, L., Qian, J., Bollen, M. & Saurin, A. T. The Importance of Kinase-Phosphatase Integration: Lessons from Mitosis. Trends in cell biology 28, 6–21, doi:10.1016/j.tcb.2017.09.005 (2018).

47 Seo, S. B. et al. Regulation of histone acetylation and transcription by INHAT, a human cellular complex containing the set oncoprotein. Cell 104, 119–130 (2001).

48 Kato, K., Okuwaki, M. & Nagata, K. Role of Template Activating Factor-I as a chaperone in linker histone dynamics. J Cell Sci 124, 3254–3265, doi:10.1242/jcs.083139 (2011).

49 Liu, L. & Eisenman, R. N. Regulation of c-Myc Protein Abundance by a Protein Phosphatase 2A-Glycogen Synthase Kinase 3β-Negative Feedback Pathway. Genes Cancer 3, 23–36, doi:10.1177/1947601912448067 (2012).

50 Network, C. G. A. R. Comprehensive molecular profiling of lung adenocarcinoma. Nature 511, 543–550, doi:10.1038/nature13385 (2014).

51 Dingar, D. et al. BioID identifies novel c-MYC interacting partners in cultured cells and xenograft tumors. J Proteomics 118, 95–111, doi:10.1016/j.jprot.2014.09.029 (2015).

52 Hsu, T. Y. et al. The spliceosome is a therapeutic vulnerability in MYC-driven cancer. Nature 525, 384–388, doi:10.1038/nature14985 (2015).

53 Futschik, M. E. & Carlisle, B. Noise-robust soft clustering of gene expression time-course data. J Bioinform Comput Biol 3, 965–988 (2005).

54 Sharma, K. et al. Ultradeep human phosphoproteome reveals a distinct regulatory nature of tyr and ser/thr-based signaling. Cell Rep 8, 1583–1594, doi:10.1016/j.celrep.2014.07.036 (2014).

55 Wu, R. et al. A large-scale method to measure absolute protein phosphorylation stoichiometries. Nat Methods 8, 677–683, doi:10.1038/nmeth.1636 (2011).

56 Pinna, L. A. & Donella-Deana, A. Phosphorylated synthetic peptides as tools for studying protein phosphatases. Biochim Biophys Acta 1222, 415–431 (1994).

57 Pinna, L. A. & Ruzzene, M. How do protein kinases recognize their substrates? Biochim Biophys Acta 1314, 191–225 (1996).

58 Hein, J. B., Hertz, E. P. T., Garvanska, D. H., Kruse, T. & Nilsson, J. Distinct kinetics of serine and threonine dephosphorylation are essential for mitosis. Nat Cell Biol 19, 1433–1440, doi:10.1038/ncb3634 (2017).

59 Schweiger, R. & Linial, M. Cooperativity within proximal phosphorylation sites is revealed from large-scale proteomics data. Biol Direct 5, 6, doi:10.1186/1745-6150-5-6 (2010).

60 Chung, V. et al. Safety, tolerability, and preliminary activity of LB-100, an inhibitor of protein phosphatase 2A, in patients with relapsed solid tumors. Clin Cancer Res, doi:10.1158/1078-0432.CCR-16-2299 (2016).

61 Oaks, J. J. et al. Antagonistic activities of the immunomodulator and PP2A-activating drug FTY720 (Fingolimod, Gilenya) in Jak2-driven hematologic malignancies. Blood 122, 1923–1934, doi:10.1182/blood-2013-03-492181 (2013).

62 Kunttas-Tatli, E., Bose, A., Kahali, B., Bishop, C. P. & Bidwai, A. P. Functional dissection of Timekeeper (Tik) implicates opposite roles for CK2 and PP2A during Drosophila neurogenesis. Genesis 47, 647–658, doi:10.1002/dvg.20543 (2009).

63 Lord, C. J. & Ashworth, A. BRCAness revisited. Nat Rev Cancer 16, 110–120, doi:10.1038/nrc.2015.21 (2016).

64 Zheng, X. F., Kalev, P. & Chowdhury, D. Emerging role of protein phosphatases changes the landscape of phospho-signaling in DNA damage response. DNA Repair (Amst) 32, 58–65, doi:10.1016/j.dnarep.2015.04.014 (2015).

65 Linding, R. et al. Systematic discovery of in vivo phosphorylation networks. Cell 129, 1415–1426, doi:10.1016/j.cell.2007.05.052 (2007).

66 Meyers, J., Craig, J. & Odde, D. J. Potential for control of signaling pathways via cell size and shape. Curr Biol 16, 1685–1693, doi:10.1016/j.cub.2006.07.056 (2006).

67 Pemovska, T. et al. Individualized systems medicine strategy to tailor treatments for patients with chemorefractory acute myeloid leukemia. Cancer discovery 3, 1416–1429, doi:10.1158/2159-8290.CD-13-0350 (2013).

68 Yadav, B. et al. Quantitative scoring of differential drug sensitivity for individually optimized anticancer therapies. Sci Rep 4, 5193, doi:10.1038/srep05193 (2014).

69 Shu, S. et al. Response and resistance to BET bromodomain inhibitors in triple-negative breast cancer. Nature 529, 413–417, doi:10.1038/nature16508 (2016).

70 Tibaldi, E. et al. Targeted activation of the SHP-1/PP2A signaling axis elicits apoptosis of chronic lymphocytic leukemia cells. Haematologica 102, 1401–1412, doi:10.3324/haematol.2016.155747 (2017).

71 Guergnon, J. et al. PP2A targeting by viral proteins: a widespread biological strategy from DNA/RNA tumor viruses to HIV-1. Biochim Biophys Acta 1812, 1498–1507, doi:10.1016/j.bbadis.2011.07.001 (2011).

72 Kauko, O. et al. PP2A inhibition is a druggable MEK inhibitor resistance mechanism in KRAS mutant lung cancer cells. Submitted for publication (2018).

73 Ritchie, M. E. et al. limma powers differential expression analyses for RNA-sequencing and microarray studies. Nucleic Acids Res 43, e47, doi:10.1093/nar/gkv007 (2015).

74 Kumar, L. & Futschik, M. Mfuzz: a software package for soft clustering of microarray data. Bioinformation 2, 5–7 (2007).

75 Hornbeck, P. V. et al. PhosphoSitePlus, 2014: mutations, PTMs and recalibrations. Nucleic Acids Res 43, D512–520, doi:10.1093/nar/gku1267 (2015).

76 Merico, D., Isserlin, R., Stueker, O., Emili, A. & Bader, G. D. Enrichment map: a network-based method for gene-set enrichment visualization and interpretation. PLoS One 5, e13984, doi:10.1371/journal.pone.0013984 (2010).

77 Shannon, P. et al. Cytoscape: a software environment for integrated models of biomolecular interaction networks. Genome research 13, 2498–2504 (2003).

78 Horn, H. et al. KinomeXplorer: an integrated platform for kinome biology studies. Nat Methods 11, 603–604, doi:10.1038/nmeth.2968 (2014).

79 Chou, M. F. & Schwartz, D. Biological sequence motif discovery using motif-x. Curr Protoc Bioinformatics Chapter 13, Unit 13.15–24, doi:10.1002/0471250953.bi1315s35 (2011).

80 Subramanian, A. et al. Gene set enrichment analysis: a knowledge-based approach for interpreting genome-wide expression profiles. Proc Natl Acad Sci U S A 102, 15545–15550, doi:10.1073/pnas.0506580102 (2005).

